# Integrative systems biology approach identified crucial genes and transcription factors associated with gallbladder cancer pathogenesis

**DOI:** 10.1101/2021.04.09.439151

**Authors:** Nabanita Roy, Mrinmoy Kshattry, Susmita Mandal, Mohit Kumar Jolly, Dhruba Kumar Bhattacharyya, Pankaj Barah

**Affiliations:** Department of Molecular Biology and Biotechnology, Tezpur University, Napaam, Sonitpur, Assam, 784028, India; Centre for BioSystems Science and Engineering, Indian Institute of Science, Bangalore-560012, India; Department of Computer Science and Engineering, Tezpur University, Napaam, Sonitpur, Assam, 784028, India

**Keywords:** Gallbladder cancer, Transcriptomics, Differentially Expressed Genes, Coexpression network, transcription factors, epithelial-mesenchymal transition, Cell cycle machinery

## Abstract

Gallbladder cancer (GBC) has a lower incidence rate among the population relative to other cancer types but majorly contributes to the total cancer cases of the biliary tract system. GBC is distinguished from other malignancies due to its high mortality, marked geographical variation and poor prognosis. To date no systemic targeted therapy is available for GBC. The main objective of this study is to determine the molecular signatures correlated with GBC development using integrative system level approaches. We performed analysis of publicly available transcriptomic data to identify differentially regulated genes and pathways. Co-expression network analysis and differential regulatory network analysis identified hub genes and hub transcription factors (TFs) associated with GBC pathogenesis and progression. We then assessed the epithelial-mesenchymal transition (EMT) status of the hub genes using a combination of three scoring methods. The hub genes such as; CDC6, MAPK15, CCNB2, BIRC7, L3MBTL1 identified are regulators of cell cycle components which suggests that cell cycle regulatory genes are significantly linked to GBC pathogenesis and progression.

## 1. Introduction

Gallbladder is a small sac like structure located beneath the liver and forms an integral component of the biliary tract system. It is the sixth most frequent cancer of the gastrointestinal tract worldwide. Gallbladder cancer (GBC) is an aggressive malignancy with rapid progression, poor prognosis and a high mortality rate characterized with an overall 5-year survival rate of 5% [1][2]The incidence rate of GBC is highly marked by distinct geographic and ethnic disparity. This regional and ethnic discrepancy in the incidence ratios of GBC cases indicates the differences in GBC etiology in different populations [3][4]. According to the Globocan report, GBC ranks at 20^th^ position among the most frequent cancer types with approximately 0.2 million cases diagnosed annually (http://globocan.iarc.fr). GBC incidence frequency is maximum in the region of Eastern Europe, East Asian countries and Latin American populations. The incidence ratio of GBC cases is the highest among South American countries such as Chile, Bolivia and Ecuador and Asian countries which mainly include Korea, India, Japan and Pakistan [5][6].

GBC is an orphan disease and its etiology is multifactorial. The pathological spectrum of GBC mainly progresses from metaplasia to dysplasia with subsequent carcinoma-in-situ and cancer metastasis which suggests that an Epithelial Mesenchymal Transition (EMT) event might be an important phenomenon in GBC development. The detailed molecular mechanism of associated risk factors in GBC is not understood yet. There is no targeted therapy available for GBC treatment. Hence, understanding of GBC pathogenesis is urgently needed for the development of targeted therapy and to improve the treatment outcome of GBC patients [7][8]. At present, the most common approach for treating GBC is radical resection. However, the majority of patients with GBC cannot undergo surgical resection due to aberrant clinical manifestations. The symptoms become noticeable in cases where the cancer has already invaded the nearby organs. In such cases, non-surgical therapies such as chemo and radiotherapy are the only options for treatment. According to the National Comprehensive Cancer Network, the single-agent therapy, which includes fluoropyrimidine or gemcitabine-based treatment, and combination therapy regimen, which includes oxaliplatin, cisplatin and capecitabine are the two chemotherapeutics option for GBC patients but both of these are still undergoing clinical trials. Till now there is no diagnostic & prognostic biomarker that can detect GBC at the initial stage to potentially select patients who are most likely to benefit from chemotherapy [9][10][11][12].

Systems biology has been responsible for some of the most important developments in the field of human health. It is a multidisciplinary approach to determine the complexity of biological systems and is used to discover new biomarkers for disease, drug targets and other treatments. The advancement of high throughput next generation sequencing (NGS) strategies in recent years such as transcriptome sequencing has helped to generate robust cancer based datasets and the analysis of these datasets using integrated system biology approaches has provided a basis for investigation of genes and their pathological functions in malignancy and its implication in cancer treatment strategies [13][14]. Weighted gene co-expression network analysis (WGCNA) is a frequently used systems biology based method for determining the gene-gene correlation across samples which can be used to identify modules containing clusters of highly correlated gene networks [15].

To this end, we have carried out transcriptomic analysis of GBC RNAseq dataset downloaded from NCBI-GEO database. We have identified potential genes and TFs associated with GBC progression and pathogenesis through co-expression network analysis on normal and GBC samples followed by regulatory network analysis. Functional enrichment analysis and EMT score calculation has also been carried to identify crucial genes for GBC pathogenesis.

## 2. Results

### 2.1 Differential gene expressions in gallbladder cancer

To identify the differential expressed molecular signature in GBC, we have carried out transcriptomic analysis in 10 tumor samples and 10 tumors’ adjacent control samples of GBC patients and subsequently visualized the normalized and transformed data using dispersion and principal component analysis [**Fig1A-B**]. Two separate clusters for GBC and control samples were identified in the PCA plot. However, the PCA plot also showed that three control samples were diverted towards the GBC cluster. This might be due to invasion of cancer cells to the adjacent control samples of GBC patients. From the transcriptomic analysis, 2980 significant DEGs were identified in GBC as compared to that of controls by taking Padj ≤ 0.05. The hierarchical clustering analysis of significant DEGs [**Fig 1C**] showed that the GBC and the adjacent control samples showed different gene expression cluster. The significant DEGs identified in GBC are largely linked to the cell cycle regulation and development processes **[Fig 1D]**. This suggests that genes related to cell cycle progression and checkpoints regulatory proteins might be crucial for GBC development.

**Fig 1:**
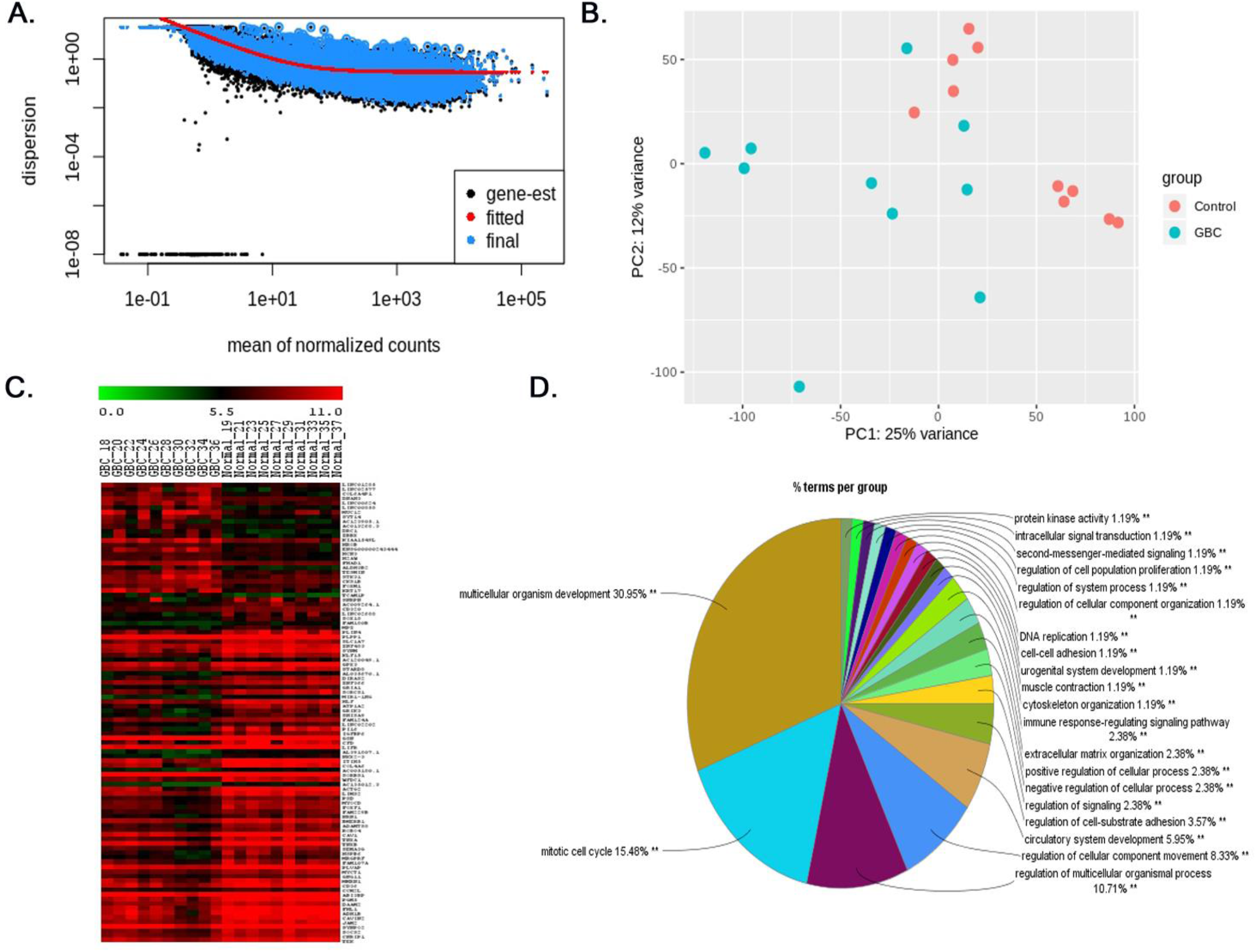
Differential gene expression in GBC. **A.** An estimate of the dispersion plot for mean of normalized counts. **B.** Principle component analysis of 10 GBC and 10 normal matxhed control samples. **C.** Hierarchical clustering of top 100 significant DEGs in GBC compared to that of control. **D.** Significant biological processes associated with GBC.

### 2.2 Construction of gene co-expression network and module detection

To identify significant DEGs module, WGCNA analysis was performed by taking 2980 significant and log2 transformed DEGs. The co-expression networks were constructed for GBC and control condition separately. The co-expression network construction needs the selection of soft thresholding power β against which the adjacency (Adj_ij_) matrix of the selected DEGs were calculated which is required to build standard scale-free coexpression network. The soft-thresholding power β for GBC and control are 18 and 20 respectively **(Supplementary Fig 1).** The Hierarchical clustering is performed for identification of modules from the constructed network of both GBC and control [**Fig 2A-B**]. The eigenmodules were clustered through dissimilarity of module eigengenes. If the correlation between the module eigenegenes are greater than 0.75, then those modules were combined into a single module. A total of 21 and 19 modules were extracted from GBC network and control network respectively. The heatmap plot for the genes in control and GBC network is represented in **Fig 2C-D**

**Fig 2:**
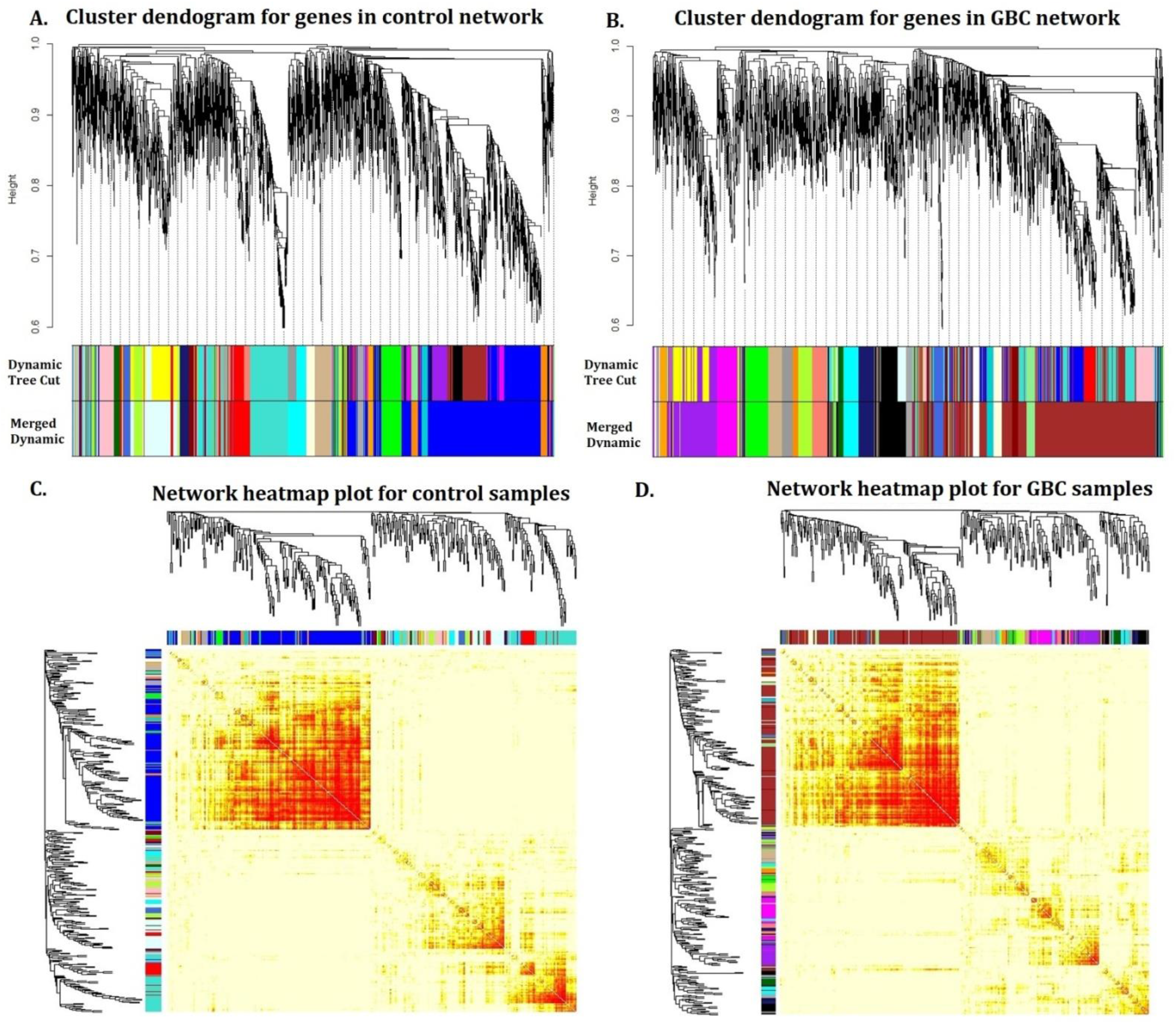
Gene co-expression network analysis: **A-B**. Clustering dendogram of genes in GBC and control network respectively. **C-D**. Clustering dendogram heatmap based on topological overlapping for GBC samples and control samples respectively.

### 2.3 Detection of nonpreserved module from GBC and control co-expression network

In this study, the module preservation analysis was performed by the following approaches: (i) GBC vs. control, where the cancer data is considered as the test data and the reference data is the control data. (ii) Control vs. GBC, in which the control data is considered as the test data and the GBC data is the reference data. The identification of nonpreserved modules in both control **[Fig 3A]** and GBC networks **[Fig 3B]** can give insights of distinct molecular signatures in GBC modules compared to that of control modules.

**Fig 3:**
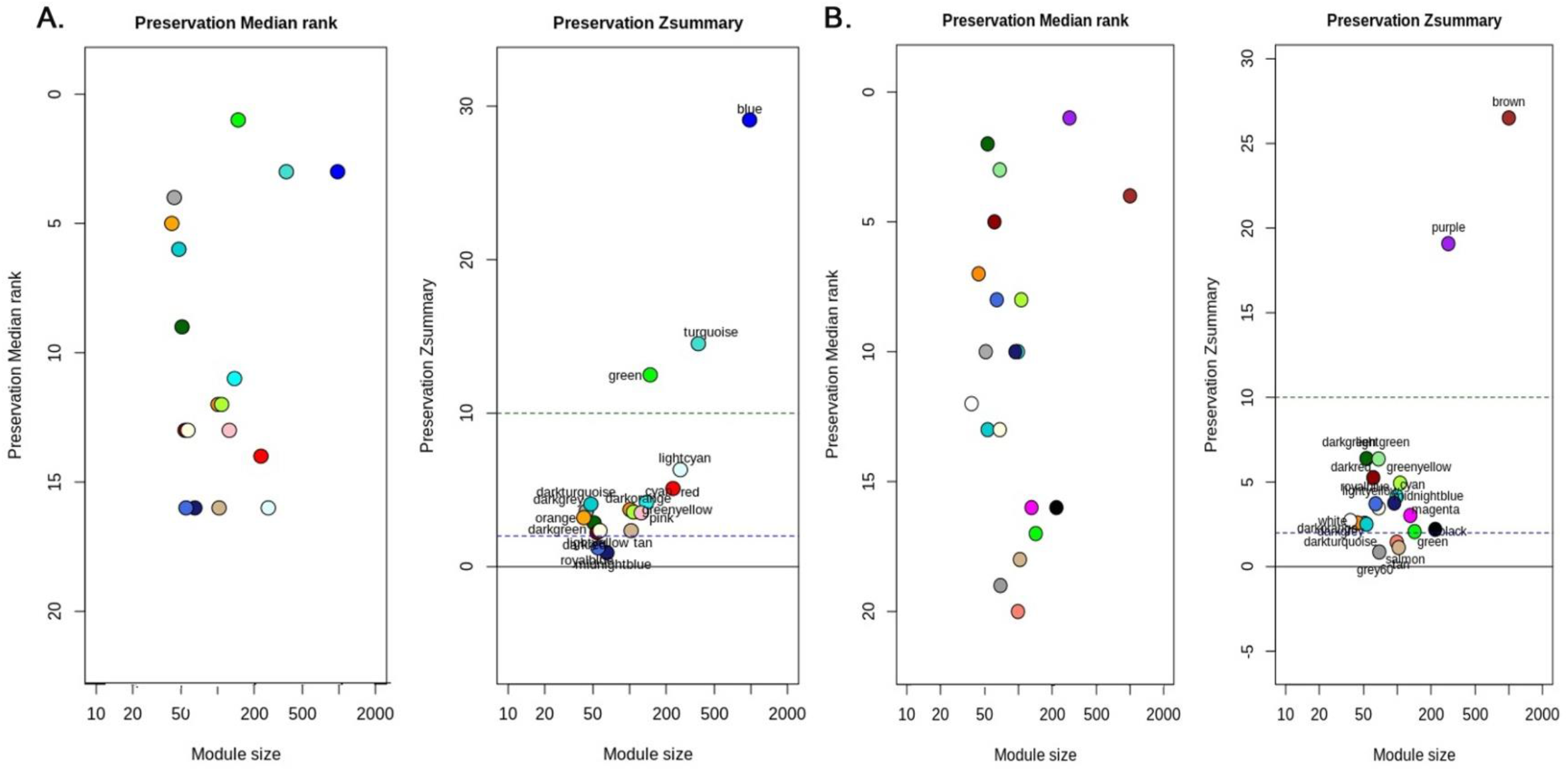
Preservation analysis of modules based on *Zsummary* and *MedianRank*. **A.** Identification of modules in the control condition. The modules in midnightblue and royalblue color is nonpreserved **B.** Identification of modules in the GBC condition. The modules in tan, salmon and grey60 color is nonpreserved.

In GBC to control module preservation analysis, three modules, salmon, tan and grey60 were identified as the non preserved module in GBC with Zsummary- 1.1, 1.4, 0.86 and median rank-20, 18, 19 respectively (**Supplementary Table1**). For control to GBC preservation analysis, two nonpreserved modules have been detected which are midnightblue and royalblue with Zsummary preservation- 0.91, 1.2. The Median Rank of both midnight and royalblue modules were 16 respectively (**Supplementary Table 2**).

### 2.4 Hub gene identification from nonpreserved modules

The genes which have high degree of connectivity or high correlations in significant modules are regarded as hub genes. The hub genes play a significant role in network biology study. For hub gene identification, we have considered the nonpreserved modules identified from both GBC and control network and determined their topological measure with respect to intra-modular connectivity. A total of five genes have been considered as potential candidate in terms of correlation weight (degree). The weight of the potential candidate genes identified through intra-modular connectivity analysis is given in (**Table 1**). The gene with highest intra-modular connectivity from each module (hub gene) were AL009178.3 (novel transcript), ADAM18 (ADAM Metallopeptidase domain 18), MAPK (Mitogen Activated Protein Kinase 15), L3MBTL1 (Lethat(3) nalignant brain tumor-like protein 1) and ALPPL2 (Alkaline phospatase, placental-like 2). Subsequently, PPI networks for each of the non-preserved modules were constructed (Fig 4) and the hub genes were identified based on degree centrality (**Table 2**). These genes are BIRC7 (Baculoviral IAP repeat-containing protein 7), CCNB2 (Cyclin B2), CDC6 (Cell division cycle 6), L3MBTL1 and WDR88 (WD repeat domain 88). All the identified hub genes are found to be upregulated in GBC as compared to controls. This indicates that upregulation of these hub genes might drive GBC development and progression.

**Table1.** Identification of gene through intra-modular connectivity analysis

| Control to GBC |  |  |  | GBC to control |  |  |  |  |  |
| --- | --- | --- | --- | --- | --- | --- | --- | --- | --- |
| midnightblue |  | royalblue |  | salmon |  | tan |  | grey60 |  |
| Gene | weight | Gene | weight | Gene | weight | Gene | weight | Gene | weight |
| AL009178.3 | 2.87 | ADAM18 | 2.87 | MAPK15 | 12.51 | L3MBTL1 | 11.38 | ALPPL2 | 8.87 |
| SPATC1 | 2.71 | CNTN4 | 2.71 | TRAPPC9 | 12.08 | ZNF337-AS1 | 10.47 | PATE4 | 8.01 |
| CTSV | 2.70 | NUP62CL | 2.70 | OPLAH | 11.72 | AC099661.1 | 9.64 | AP00842.3 | 7.85 |
| AL353746.1 | 2.64 | QTRT2 | 2.64 | OTUD6B | 10.97 | AC240565 | 8.68 | GPATCH1 | 7.80 |
| AL360270.1 | 2.61 | LINCO1517 | 2.61 | TAF2 | 10.72 | C1QTNF | 8.14 | AP000977.1 | 7.06 |

**Fig 4:**
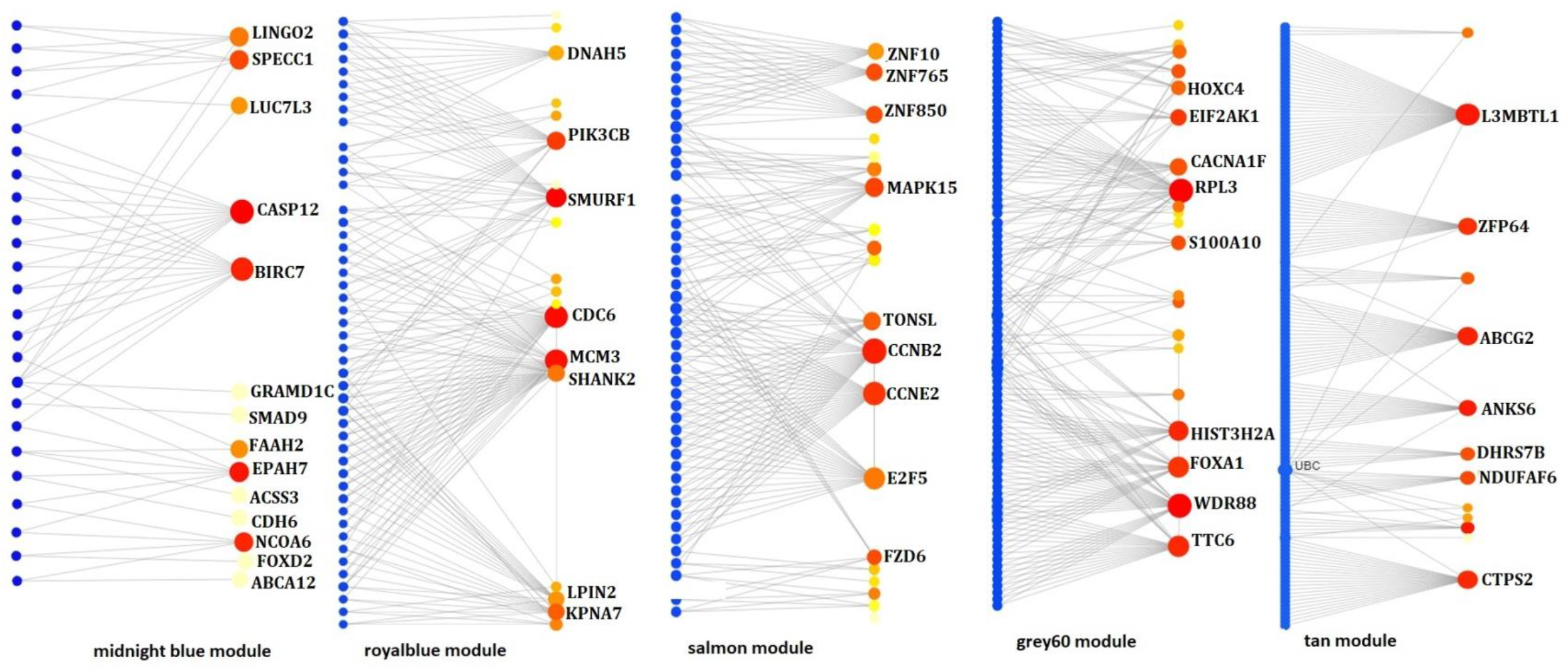
PPI network analysis of nonpreserved modules. The small blue circles represent the proteins and large red node represents the genes in the modules

**Table2.**
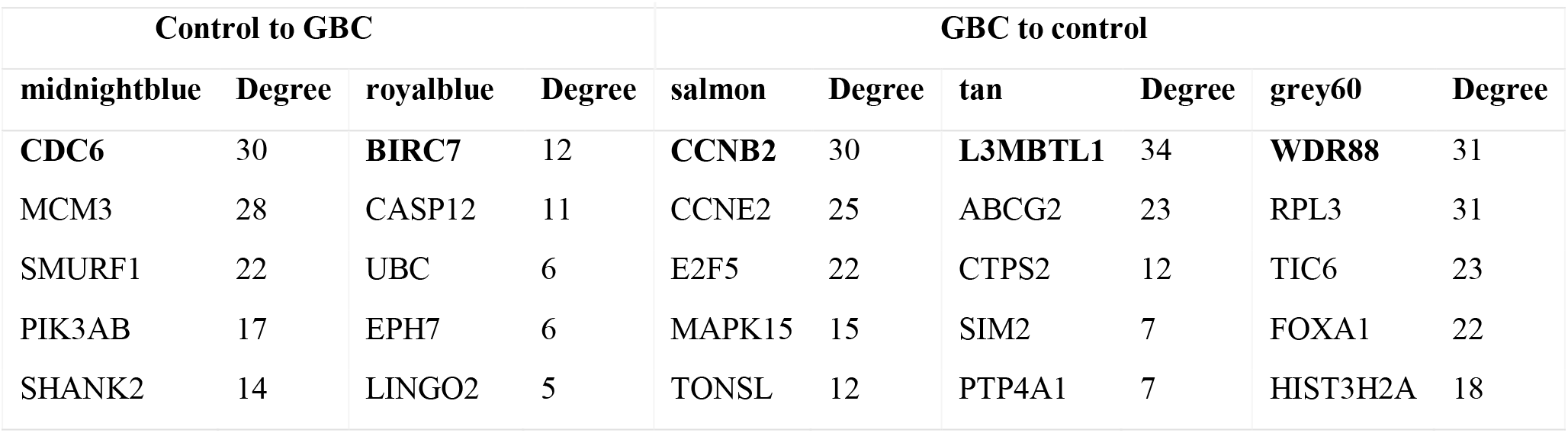
Hub gene identification through PPI network analysis

### 2.5 Functional and annotation and pathway associated with genes of the nonpreserved modules

The functional GO terms and pathways associated with the gene modules were identified using DAVID. The statistical significance of p-value<0.05 were considered for finding important biological processes and KEGG pathways related to GBC progression. The functional annotation analysis identified that the module genes were mainly associated with cell cycle regulation processes, metabolic pathways and signal transduction processes. The top ranked significant biological processes and pathways were tabulated in **Table 3A** and **3B** respectively

**Table 3A.**
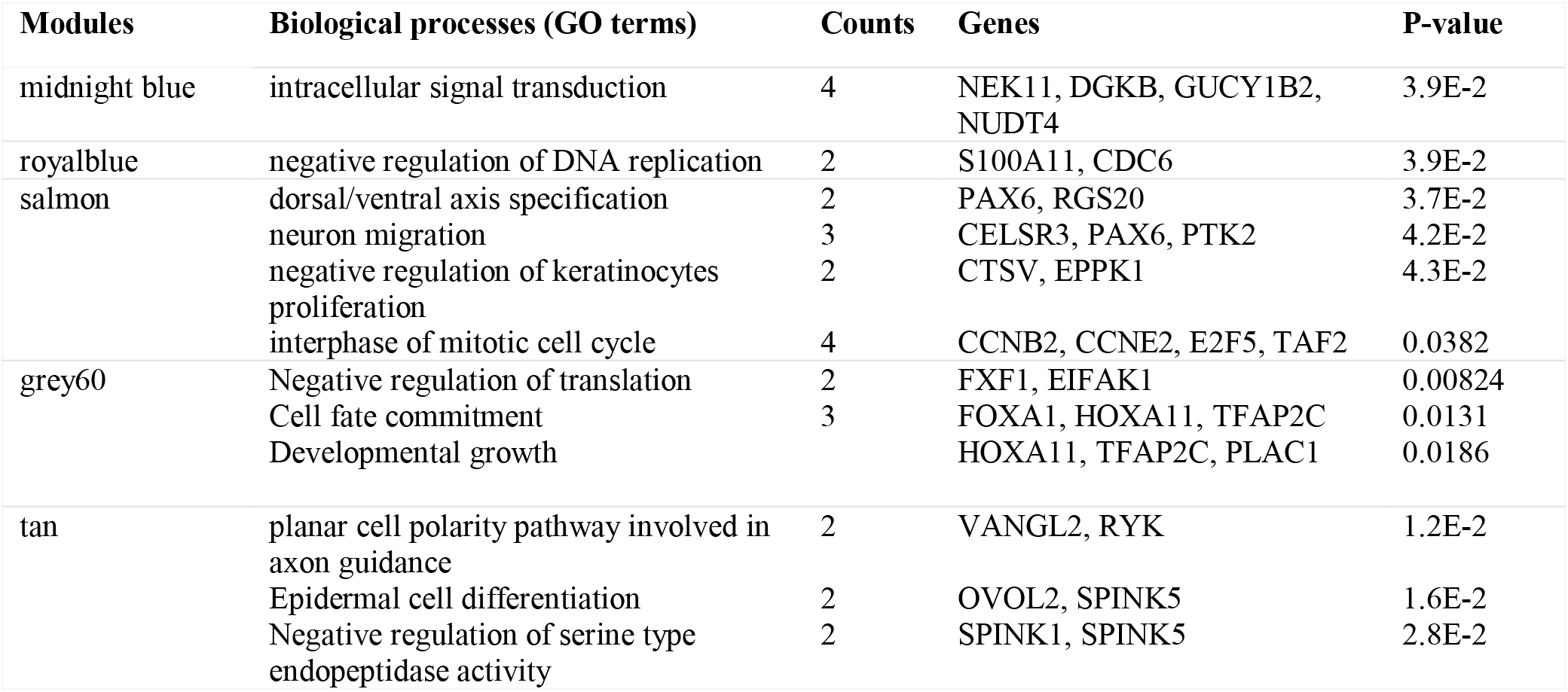
Enriched biological processes associated with nonpreserved modules identified from GBC network

**Table 3B.** Enriched KEGG pathways associated with nonpreserved modules identified from GBC network

| Module | KEGG pathways | counts | genes | p-value |
| --- | --- | --- | --- | --- |
| midnightblue | Glycerolipid metabolism | 2 | HLA-DMA | 0.00683 |
|  | Toxoplasmosis | 2 | DGKB, LIPC | 0.0222 |
|  | Apoptosis | 2 | CTSV, CASP12 | 0.0313 |
|  | Glycosaminoglycan degradation | 1 | HYAL4 | 0.0382 |
| royalblue | Phosphatidylinositol signaling | 2 | PIK3CB, ITPKA | 0.00762 |
|  | Cell cycle | 2 | MCM3, CDC6 | 0.0204 |
| salmon | Cell cycle | 3 | E2F5, CCNB2, CCNE2 | 4.5E-2 |
|  | Small cell lung cancer | 2 | CCNE2, PTK2 | 0.0385 |
|  | P53 signaling | 2 | CCNE2, CCEB2 | 0.0240 |
|  | Ubiquitin biosynthesis | 1 | COQ2T | 0.0364 |
| grey60 | Thiamine metabolism | 1 | ALPPL2 | 0.0266 |
|  | Necroptosis | 2 | H2AW, RNF103-CHMP3 | 0.0292 |
|  | Alcoholism | 2 | H2AW, H2BO1 | 0.0355 |
|  | Histidine metabolism | 1 | ALDH3B2 | 0.0380 |
| tan | Wnt signaling pathway | 2 | VANGL2, RYK | 0.0321 |
|  | Steroid biosynthesis |  | CYP24A1 | 0.0339 |

### 2.6 Identification of hub transcription factors in GBC through TG-TF regulatory network analysis

Out of the 2980 DEGs, 106 of the DEGs code for transcription factors (TFs). Considering these TFs as the source nodes and the DEGs (including the TFs) as the target nodes, we created a transcription network. The degree distribution did not follow a Poisson distribution (mean of degree distribution = 78.88603; variance of degree distribution = 41142.97) and hence, the network is not a random network. The topological parameters of the network such as clustering coefficient, path length, assortavity were calculated using the R package igraph. The assortavity degree of the network is negative i.e. −0.1024318, meaning the nodes with higher degrees tend to interact with nodes of smaller degrees. This is in compliance with the observation that real-world networks tend to have negative assortavity. The degree coefficient,Ɣ, of the degree distribution was calculated to be 5.467 and a power-law was fitted in the distribution The hub TFs identified in GBC were PAX6, KLF15, NR2F1, TFAP2C, FOXJ2 and FLR **[Fig 5A]** Among these, PAX6, TFAP2C and FOXJ2 were present in modules identified from GBC co-expression network.

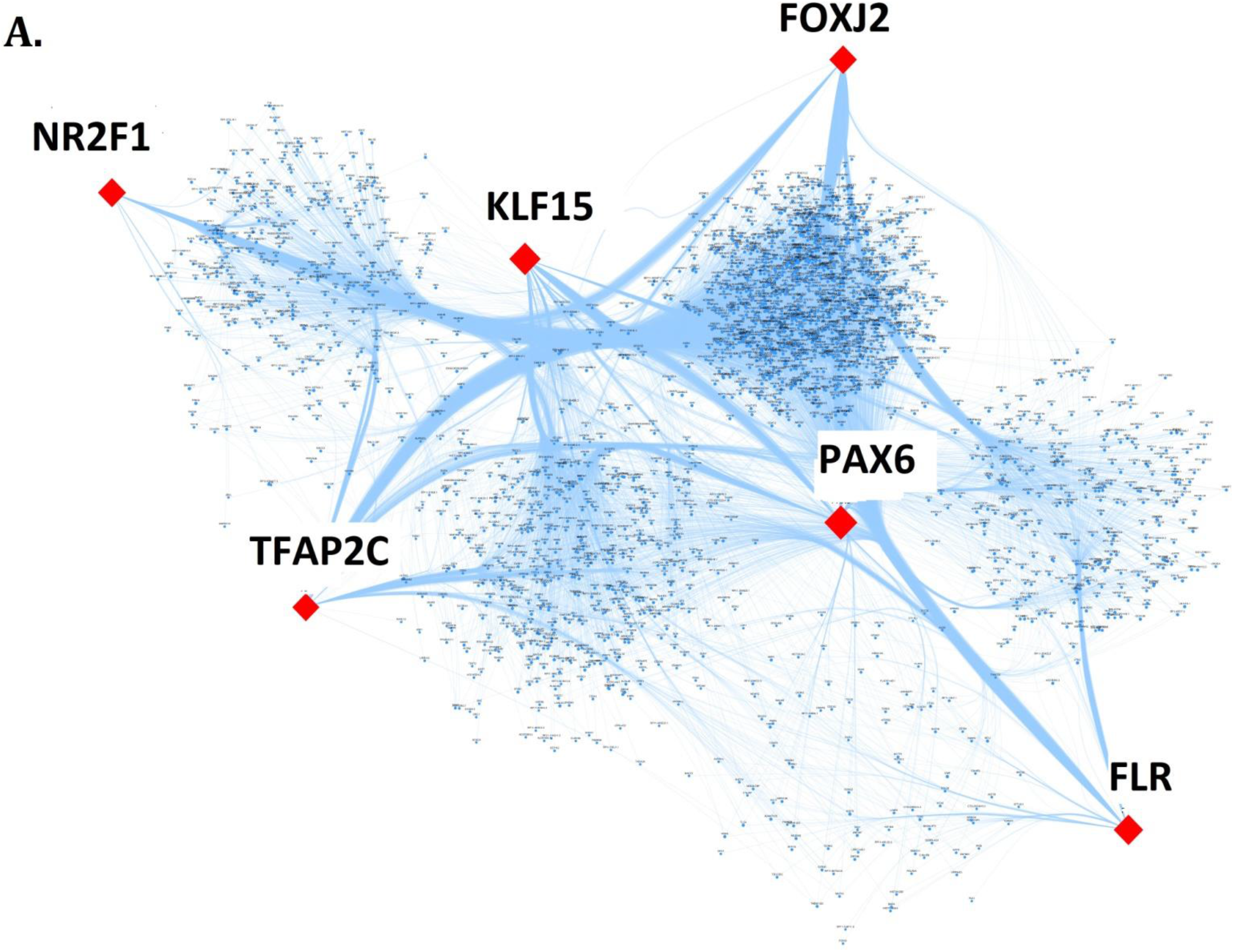

**Fig 5:**
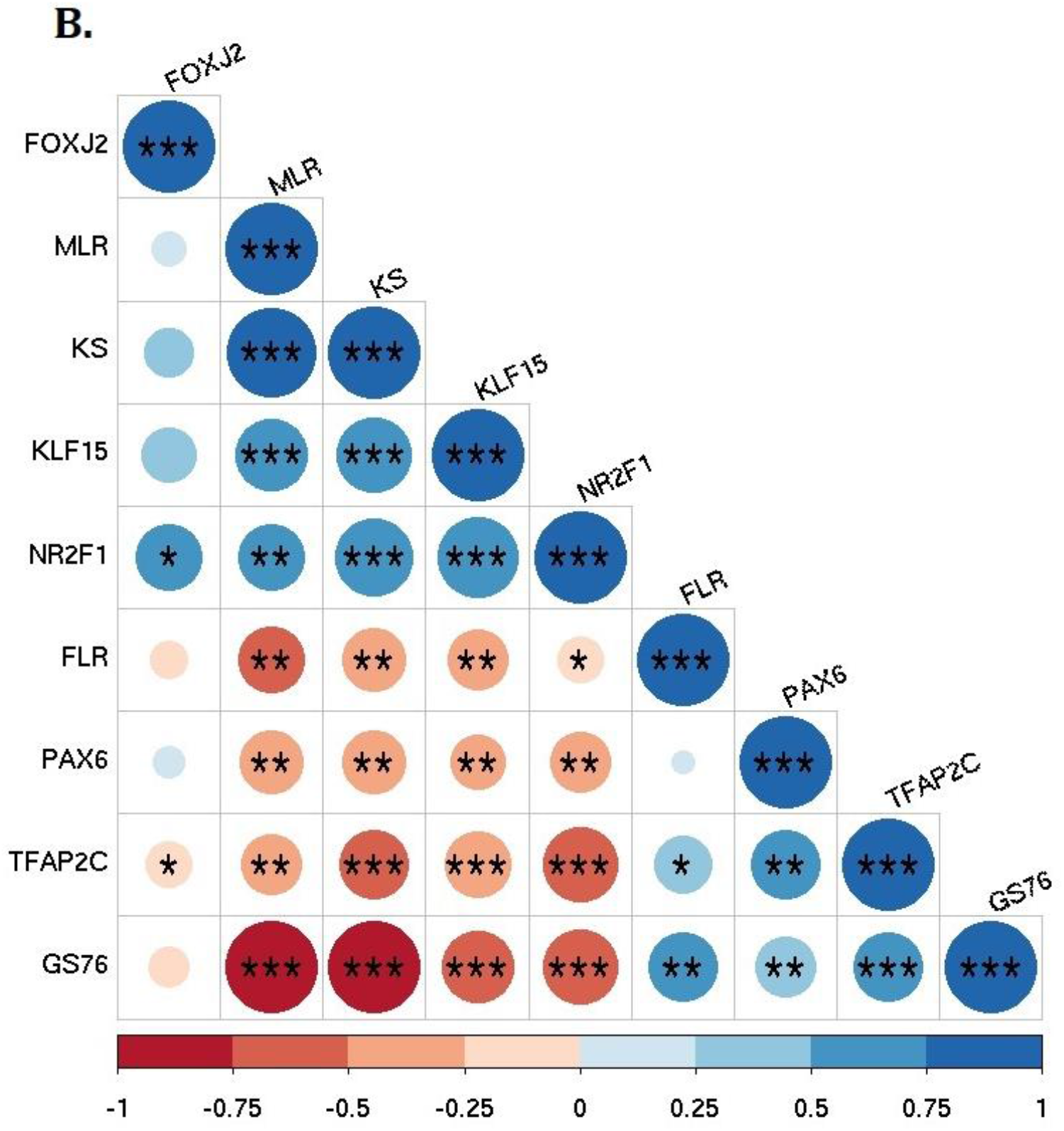
Regulatory network analysis of DEGs in GBC. **A.** Identification of hub TFs in GBC. The red node represents the top hub TFs and the small blue node represents target genes. **B.** Pairwise correlation of hub transcription factor identified through TG-TF interaction.

### 2.7 EMT analysis identified differential EMT patterns in hub Transcription Factors

Next, we quantified the extent of epithelial-mesenchymal transition (EMT) that the GBC samples have gone through. We used three different algorithms (76GS, KS, MLR) that score the degree of EMT in transcriptomic data – while higher KS and MLR scores indicate a more mesenchymal states, a higher 76GS score indicates a more epithelial phenotype (**Supplementary Table 3**). Thus, 76GS scores are expected to correlate negatively with KS and MLR scores for GBC samples, based on previous observations [16]. Indeed we observed a positive significant correlation between MLR and KS scores and both the scores are negatively correlated with 76GS scores. This consistency indicates that these EMT scoring methods well recapitulate the extent of EMT in GBC samples. Further, we examined how the six hub TFs identified in GBC were associated with coordinating a more epithelial vs. a more mesenchymal phenotype. Levels of KLF15 and NR2F1 associated with a more mesenchymal state (i.e. positive correlation with KS and MLR scores and negative correlation with 76GS scores). FOXJ2 also showed similar trends but they were not statistically significant. On the other hand, FLR, PAX6 and TFAP2C were associated with an epithelial state (i.e. positive correlation with 76GS scores, and negative correlation with KS and MLR scores). Thus, the six hub TFs identified in GBC associate differentially with epithelial vs. mesenchymal status, seemingly forming two ‘teams’ of players – one promoting EMT, the other set inhibiting EMT **[Fig 5B]**.

## 3. Discussions

Gallbladder cancer is a fatal malignancy of the biliary tract system. Standard molecular screening of GBC is the utmost necessity to detect the onset of GBC at an early stage and to reduce mortality rate of patients suffering from GBC. The use of next generation sequencing techniques is being widely used in cancer related studies. However, limited integrative omics based studies have been carried out in GBC due to the rarity of the disease. The therapeutic strategies of GBC are also limited due to lack of specific molecular targets. Hence, the main objective of this study was to identify important genes and TFs related to GBC pathogenesis.

To this end, we have carried out system based approach to screen hub genes/systems biomarker in GBC. Here we have analyzed transcriptomic dataset consisting of 10 GBC and 10 adjacent control samples. In total 2980 significant DEGs were identified in GBC samples compared to that of the controls. The identified DEGs were then used to construct gene co-expression network and to determine significant co-expressed modules in both GBC and control samples. The functional annotations and KEGG pathway analysis were further evaluated to identify significant biological processes and pathways enriched in nonpreserved modules. We analyzed and identified the hub transcription factors from significant DEGs that might have important role in gene regulation process during GBC development. The hub genes identified from the nonpreserved coexpressed modules were largely associated with cell cycle machinery and signaling processes.

The cell cycle machinery is a highly regulated and intricate process which governs the cell growth, cell proliferation and cell division through its cell cycle regulatory genes. The cell cycle regulatory molecule majorly involves growth-regulatory signaling proteins- CDK and CDKI and associated genes/proteins that checks for any anomalies throughout the genome. Disruption in the regulation of cell cycle machinery/components are frequently observed in several malignancies where it contributes to malignant transformation and resistant to cancer drugs [17, 18]. Numerous studies in the last two decades have reported the significance of cell cycle aberration towards human cancer development. The cell cycle defects in cancer mainly involves uncontrolled proliferation through dysregulation in any of its cell cycle components either due to CDK function misregulation and/or decrease in the negative regulator of CDKI [19, 20]. The most important component of cell cycle machinery is the DNA replication initiation process and pathway. The DNA synthesis process acts as a relay system of the cell cycle process the connects various growth signaling network with DNA replication complex and therefore this component serves as an important diagnostic and prognostic target[21]. The DNA replication and the mitotic process regulation are considered to be the central players involved in these cell cycle phase transitions and therefore they are not only as useful cancer biomarkers but also as potent targets for mechanism based therapies [22]. But the initiation of oncogenesis process is not only associated with cell cycle components alone. The development of malignant tumors involves mis-regulation of the cell death machinery and cell–cell and/or cell–matrix interactions that co-operate with cell cycle defects[23].

The hub genes identified through gene co-expression network analysis followed by PPIs analysis were directly or indirectly associated with components of the cell cycle system, apoptotic regulation and cell-cell adhesion process that ultimate give rise to uncontrolled cell proliferation and later on to a full bloom malignancy. The hub genes L3MBTL1, MAPK15, CCNB2 and CDC6 are crucial element that acts as a control system for coordinated regulation of cell cycle system. The L3MBTL1 is known to be a potential tumor suppressor gene in drosophila fly. It binds to the chromatin complex during S-phase of the cell cycle also regulates the target genes of E2F-RB negatively that are necessary for S-phase initiation. L3MBTL1 was reported to be associated with breast cancer and myeloid leukemia including AML [24–26]. The family of MAPK proteins plays a key role in different cellular events such as cell differentiation, cell growth and development, cellular transformation and apoptosis. It involves a sequence of protein kinase signaling cascade which is important for regulation of cellular proliferation[27]. MAPK15 is known to be an important extracellular signal transducing kinase which is known to be activated by human serum. The MAPK15 gene is unique as it does not have specific MEK upstream regulation like other MAP kinases. The activity of MAPK15 is found to be modulated by several oncogenes and recent study reported the association of MAPK15 with BCR-ABL mediated autophagy and functions in oncogene dependent cancer cell proliferation and progression[28, 29]. CCBN2 has been found to be linked with poor survival outcome in gastric and hepatocellular cancer[30, 31]. CDC6 also acts as an crucial player in cell cycle system and acts as a replication licensing factor and governs the DNA replication process through maintenance of the cell cycle checkpoints machinery. CDC6 is found to be reported in initial stages of many cancers and also contributes to the oncogenic activities in tumor development[32, 33].Aberrant CDC6 expression is reported to be associated with several malignancies [34]

The ADAM18, a membrane anchored gene (matrix metalloproteinase) of the ADAM family proteins that regulates cell adhesion via their interaction with integrins. It plays an important role in the release of biologically important ligands, such as tumor necrosis factor-alpha, epidermal growth factor, transforming growth factor-alpha, and amphiregulin[35].In human cancer, overexpression of specific ADAMs is related to tumor progression and poor outcome. It is regarded as a potential targets for the cancer therapeutics, particularly those cancers that are n human epidermal growth factor receptor (EGFR) ligands or TNF-alpha dependent [36, 37].

ALPPL2 belongs to the member of ALPP alkaline phosphate which is reported to be associated with tumor initiation. It was reported as a specific and targeted tumor cell surface antigen. It is significantly associated with gastric cancer and pancreatic cancer and also acts as a novel protein in pancreatic cancer[38, 39].

BIRC7 is a novel member of the IAP family and is found to be highly overexpressed in various cancer types. BIRC7 was found to be overexpressed in 66% of the cancers and remain absent in normal cells/tissues. The function of BIRC7 gene is mainly related to apoptotic regulation and signaling process. The overexpression of BIRC7 in cancers is reported to be associated with cancer drug and radio resistances, disease recurrence and poor survival[40, 41]. Moreover, increased expression of BIRC7 was found in extrahepatic cholangiocarcinoma and was significantly associated with poor prognosis and overall survival of the patient[42].WDR88 (WD repeat domain 88) present on chromosome 19 is known to be important biomarker for early prostate cancer development. This gene is evolutionary conserved and can found in 167 organisms as an ortholologus genes. Hence this gene might act as an important target in GBC [43].

We observed that genes related to cellular processes were essential and significant in pathogenesis of GBC. The genes and TFs identified from the nonpreserved modules may play key roles in the pathogenesis of GBC. The identified hub genes provided the basis for further in depth studies. In summary, this study used co-expression network analysis and transcriptional regulatory network analysis to identify key hub genes associated with GBC pathogenesis.

## 4. Materials and methods

### 4.1 Retrieval of GBC RNA-seq dataset

A comprehensive and thorough search was conducted in NCBI database for relevant dataset on GBC. The datasets were checked carefully to ascertain whether it can be considered for our study based on the following criteria- (i) the dataset must include case-control studies, (ii) the dataset must be paired end and (iii) the sequencing platform for generating the data and experimental protocol should be described in details. Based on the above mentioned criteria, we found GSE139682 from NCBI-GEO database (GEO). The dataset includes 20 samples in total obtained through resection surgery (10 samples of GBC tissues and 10 normal matched tissue samples). The GBC dataset was downloaded from NCBI-GEO database in the SRA format. The SRA files were converted to fastq using fastq-dump [http://ncbi.github.io/sra-tools]. Quality check of the fastq files was done using fastQC. The reads after quality control were aligned using Hisat2 [44] against the reference human genome *Homo sapiens* (GRCh38). The mapped reads were quantified at the gene level to obtain the count matrix for each gene using featureCounts [45]. DESeq2[46] was used for identifying differentially expressed genes (DEGs). The GBC count data was normalized and transformed in DESeq2. The level of shrinkage of each gene and the overall covariates has been estimated using dispersion plot (Fig 1A) and principle component analysis (Fig 1B) respectively. Finally, the significant DEGs of GBC were sorted by considering p-adjusted value<0.05.

### 4.2 Differential Gene Co-expression network analysis

The gene co-expression network gives a cluster of genes that are highly correlated. In comparison to other biological network analysis, differential co-expression networks can be used to build cancer specific sub-network[47]. The significant DEGs were used as input to build gene co-expression network using an R package WGCNA[15]. Using WGCNA package, two weighted gene co-expression network were constructed for cancer and control condition. For each cancer and control dataset, Pearson's correlations analysis of each gene pair was used to build an adjacency matrix using the adjacency function of the WGCNA package. Subsequently, the adjacency matrix was used to create a scale-free co-expression network based on a soft-thresholding parameter βeta (β) that is used to enrich strong correlations between gene pairs [48]. The calculated adjacency matrix was converted into Topological Overlap Matrix (TOM) by using the function TOMsimilarity. This topological overlap matrix was then used as an input for performing hierarchical clustering using the flashClust function for module identification. Finally, the network modules for cancer and control dataset were identified using dynamicTreeCut (an R package) with a minimum module size (minClusterSize = 30) and minimum sensitivity (deepSplit=2) for the gene dendrogram.

### 4.3 Module preservation analysis

The preservation analysis was performed to assess the nonpreserved module between the cancer network and control network. The basic statistics behind the module preservation is to evaluate the preservation of genes within-module between (GBC and control) a test network and reference network [15]. It has been assumed that the genes embodied in nonpreserved modules in cancer network might play a role in the pathogenesis process as compared to control network. The preservation analysis was carried out using the WGCNA function modulePreservation to determine the connectivity and weight of each module of cancer and control networks. The Preservation analysis statistics- *Z-summary* and *medianRank* gives overall significance of the preservation of a module based on degree and connectivity. The *Z-summary preservation*<2 indicates no preservation, 2≤*Z-summary*≤10 have weak to moderate preservation, and *Z-summary preservation*>10 means strong preservation [15]

### 4.4 Gene ontology and pathway analysis of the nonpreserved module

For interpreting the biological role of significant DEGs identified from nonpreserved modules, functional enrichment and pathway analysis were performed using DAVID [49].. The significant DEGs for GBC were used for the GO analysis and KEGG pathway analysis for identification of important cellular processes and pathways in GBC. The top five GO terms for biological processes and KEGG pathway terms were estimated with p-value less than 0.05.

### 4.5 Screening of hub genes from nonpreserved modules through Intramodular connectivity and PPIs network analysis

In network concept, connectivity is generally considered as the degree. In this study, we have used the intramodular connectivity approach for the screening of hub genes within weakly preserved modules. The intramodular connectivity basically measures the degree of each gene within a module. The criteria of this approach are to calculate the connectivity from the whole network (kTotal) and the connectivity within modules (kWithin). This measure of connectivity is useful to determine the biologically significant modules by calculating the degree of genes within modules. STRING database (version 10.0)[50] was used to predict potential interactions between gene candidates at the protein level. A combined score of > 0.4 was considered significant and the PPI networks of the nonpreserved modules were created using the Network analyst tool [51]. The genes with high number of connections with other genes/proteins were considered as hub gene.

### 4.6 Gene Regulatory Network (GRN) analysis

From the gene regulatory network (GRN) the information regarding the regulatory interactions between regulators and their target genes can be obtained [52]. Transcription factors are the key players in regulatory network interaction as they influence the gene expression by binding to the start site of the gene promoter region. We have used the significant DEGs as input to construct the regulatory network. The human TFs and their position weight matrices (PWMs) were downloaded from the cis-BP database [53].The matrix scan was used to predict the interaction between the TFs and its target genes. The results of the matrix-scan were filtered by setting a p-value cut off 10^−4^ The TG-TF interactions data along with their prediction scores were represented in the form of interactive network using Cytoscape software [54] (Fig. 5). Later, we have selected the top six unique hub genes considering the highest degree centrality.

### 4.7 EMT scores calculation

In this study, we have quantified the Epithelial-Mesenchymal Transition (EMT) scores for each sample using three different scoring metrics- 76Gs, MLR and KS [55–57]

## Acknowledgement

Dr. Pankaj Barah would like to acknowledge Department of Biotechnology, Government of India for providing the Ramalingaswami Re-entry Fellowship grant.

## Conflicts of interest

The authors of this manuscript have no conflict of interest

## Dataset availability

The dataset used for this study is available at NCBI-GEO database (GEO Accession No. GSE139682)

## Notes

### Competing Interest Statement

The authors have declared no competing interest.

## References

1. Wistuba II, Gazdar AF (2004) Gallbladder cancer: Lessons from a rare tumour. Nat Rev Cancer 4:695–706. https://doi.org/10.1038/nrc1429

2. Hundal R, Shaffer EA (2014) Gallbladder cancer: Epidemiology and outcome. Clin Epidemiol 6:99–109. https://doi.org/10.2147/CLEP.S37357

3. Lazcano-Ponce EC, Miquel JF, Munoz N, et al (2001) Epidemiology and Molecular Pathology of Gallbladder Cancer. CA Cancer J Clin 51:349–364. https://doi.org/10.3322/canjclin.51.6.349

4. Randi G, Franceschi S, La Vecchia C (2006) Gallbladder cancer worldwide: Geographical distribution and risk factors. Int J Cancer 118:1591–1602. https://doi.org/10.1002/ijc.21683

5. Rawla P, Sunkara T, Thandra KC, Barsouk A (2019) Epidemiology of gallbladder cancer. Clin Exp Hepatol 5:93–102. https://doi.org/10.5114/ceh.2019.85166

6. Pandey A, Stawiski EW, Durinck S, et al (2020) Integrated genomic analysis reveals mutated *ELF3* as a potential gallbladder cancer vaccine candidate. Nat Commun 1–13. https://doi.org/10.1038/s41467-020-17880-4

7. Nemunaitis JM, Brown-Glabeman U, Soares H, et al (2018) Gallbladder cancer: Review of a rare orphan gastrointestinal cancer with a focus on populations of New Mexico. BMC Cancer 18:1–14. https://doi.org/10.1186/s12885-018-4575-3

8. Bizama C, García P, Espinoza JA, et al (2015) Targeting specific molecular pathways holds promise for advanced gallbladder cancer therapy. Cancer Treat Rev 41:222–234. https://doi.org/10.1016/j.ctrv.2015.01.003

9. Song X, Hu Y, Li Y, et al (2020) Overview of current targeted therapy in gallbladder cancer. Signal Transduct Target Ther 5:. https://doi.org/10.1038/s41392-020-00324-2

10. Ben-Josef E, Guthrie KA, El-Khoueiry AB, et al (2015) SWOG S0809: A phase II intergroup trial of adjuvant capecitabine and gemcitabine followed by radiotherapy and concurrent capecitabine in extrahepatic cholangiocarcinoma and gallbladder carcinoma. J Clin Oncol 33:2617–2622. https://doi.org/10.1200/JCO.2014.60.2219

11. Weigt J, Malfertheiner P (2010) Cisplatin plus gemcitabine versus gemcitabine for biliary tract cancer. Expert Rev Gastroenterol Hepatol 4:395–397. https://doi.org/10.1586/egh.10.45

12. Ebata T, Ercolani G, Alvaro D, et al (2017) Current status on cholangiocarcinoma and gallbladder cancer. Liver Cancer 6:59–65. https://doi.org/10.1159/000449493

13. Manzoni C, Kia DA, Vandrovcova J, et al (2018) Genome, transcriptome and proteome: The rise of omics data and their integration in biomedical sciences. Brief Bioinform 19:286–302. https://doi.org/10.1093/BIB/BBW114

14. Werner HMJ, Mills GB, Ram PT (2014) Cancer systems biology: A peek into the future of patient care? Nat Rev Clin Oncol 11:167–176. https://doi.org/10.1038/nrclinonc.2014.6

15. Langfelder P, Horvath S (2008) WGCNA: An R package for weighted correlation network analysis. BMC Bioinformatics 9:. https://doi.org/10.1186/1471-2105-9-559

16. Chakraborty P, George JT, Tripathi S, et al (2020) Comparative Study of Transcriptomics-Based Scoring Metrics for the Epithelial-Hybrid-Mesenchymal Spectrum. Front Bioeng Biotechnol 8:1–13. https://doi.org/10.3389/fbioe.2020.00220

17. Otto T, Sicinski P (2017) Cell cycle proteins as promising targets in cancer therapy. Nat Rev Cancer 17:93–115. https://doi.org/10.1038/nrc.2016.138

18. Schwartz GK, Shah MA (2005) Targeting the cell cycle: A new approach to cancer therapy. J Clin Oncol 23:9408–9421. https://doi.org/10.1200/JCO.2005.01.5594

19. Malumbres M, Barbacid M (2009) Cell cycle, CDKs and cancer: A changing paradigm. Nat Rev Cancer 9:153–166. https://doi.org/10.1038/nrc2602

20. Carnero A (2002) Targeting the cell cycle for cancer therapy. Br J Cancer 87:129–133. https://doi.org/10.1038/sj.bjc.6600458

21. Williams GH, Stoeber K (2012) The cell cycle and cancer. J Pathol 226:352–364. https://doi.org/10.1002/path.3022

22. Williams GH, Stoeber K (2007) Cell cycle markers in clinical oncology. Curr Opin Cell Biol 19:672–679. https://doi.org/10.1016/j.ceb.2007.10.005

23. Bartek J, Lukas J, Bartkova J (1999) Perspective: Defects in cell cycle control and cancer. J Pathol 187:95–99. https://doi.org/10.1002/(sici)1096-9896(199901)187:1<95::aid-path249>3.0.co;2-%23

24. Addou-Klouche L, Adélaïde J, Finetti P, et al (2010) Loss, mutation and deregulation of L3MBTL4 in breast cancers. Mol Cancer 9:1–13. https://doi.org/10.1186/1476-4598-9-213

25. Sauvageau M, Sauvageau G (2010) Polycomb group proteins: Multi-faceted regulators of somatic stem cells and cancer. Cell Stem Cell 7:299–313. https://doi.org/10.1016/j.stem.2010.08.002

26. Gurvich N, Perna F, Farina A, et al (2010) L3MBTL1 polycomb protein, a candidate tumor suppressor in del(20q12) myeloid disorders, is essential for genome stability. Proc Natl Acad Sci U S A 107:22552–22557. https://doi.org/10.1073/pnas.1017092108

27. Wei Z, Liu HT (2002) MAPK signal pathways in the regulation of cell proliferation in mammalian cells. Cell Res 12:9–18. https://doi.org/10.1038/sj.cr.7290105

28. Jin DH, Lee J, Kim KM, et al (2015) Overexpression of MAPK15 in gastric cancer is associated with copy number gain and contributes to the stability of c-Jun. Oncotarget 6:20190–20203. https://doi.org/10.18632/oncotarget.4171

29. Colecchia D, Rossi M, Sasdelli F, et al (2015) MAPK15 mediates BCR-ABL1-induced autophagy and regulates oncogene-dependent cell proliferation and tumor formation. Autophagy 11:1790–1802. https://doi.org/10.1080/15548627.2015.1084454

30. Zhang HP, Li SY, Wang JP, Lin J (2018) Clinical significance and biological roles of cyclins in gastric cancer. Onco Targets Ther 11:6673–6685. https://doi.org/10.2147/OTT.S171716

31. Li R, Jiang X, Zhang Y, et al (2019) Cyclin B2 Overexpression in Human Hepatocellular Carcinoma is Associated with Poor Prognosis. Arch Med Res 50:10–17. https://doi.org/10.1016/j.arcmed.2019.03.003

32. Deng Y, Jiang L, Wang Y, et al (2016) High expression of CDC6 is associated with accelerated cell proliferation and poor prognosis of epithelial ovarian cancer. Pathol Res Pract 212:239–246. https://doi.org/10.1016/j.prp.2015.09.014

33. Lim N, Townsend PA (2020) Cdc6 as a novel target in cancer: Oncogenic potential, senescence and subcellular localisation. Int J Cancer 147:1528–1534. https://doi.org/10.1002/ijc.32900

34. Youn Y, Lee J chan, Kim J, et al (2020) Cdc6 disruption leads to centrosome abnormalities and chromosome instability in pancreatic cancer cells. Sci Rep 10:1–11. https://doi.org/10.1038/s41598-020-73474-6

35. Brocker CN, Vasiliou V, Nebert DW (2009) Evolutionary divergence and functions of the ADAM and ADAMTS gene families. Hum Genomics 4:43–55. https://doi.org/10.1186/1479-7364-4-1-43

36. Duffy MJ, McKiernan E, O’Donovan N, McGowan PM (2009) Role of ADAMs in cancer formation and progression. Clin Cancer Res 15:1140–1144. https://doi.org/10.1158/1078-0432.CCR-08-1585

37. Mullooly M, McGowan PM, Crown J, Duffy MJ (2016) The ADAMs family of proteases as targets for the treatment of cancer. Cancer Biol Ther 17:870–880. https://doi.org/10.1080/15384047.2016.1177684

38. Su Y, Zhang X, Bidlingmaier S, et al (2020) ALPPL2 Is a Highly Specific and Targetable Tumor Cell Surface Antigen. Cancer Res 80:4552–4564. https://doi.org/10.1158/0008-5472.can-20-1418

39. Dua P, Kang HS, Hong S, et al (2013) Alkaline Phosphatase ALPPL-2 Is a Novel Pancreatic Carcinoma-Associated Protein. 73:1934–1946. https://doi.org/10.1158/0008-5472.CAN-12-3682

40. Liu K, Yu Q, Li H, et al (2020) BIRC7 promotes epithelial-mesenchymal transition and metastasis in papillary thyroid carcinoma through restraining autophagy. 10:78–94

41. Liang J, Zhao W, Tong P, et al (2020) Comprehensive molecular characterization of inhibitors of apoptosis proteins (IAPs) for therapeutic targeting in cancer. 1–13

42. Li J, Yang Z, Huang S (2020) BIRC7 and STC2 Expression Are Associated With Tumorigenesis and Poor Outcome in Extrahepatic Cholangiocarcinoma. 19:1–8. https://doi.org/10.1177/1533033820971676

43. Jaiswal R, Jauhari S, Islamia JM (2017) WDR88, CCDC11, and ARPP21 genes indulge profoundly in the desmoplastic retort to prostate and breast cancer metastasis.

44. Kim D, Langmead B, Salzberg SL (2015) HISAT: A fast spliced aligner with low memory requirements. Nat Methods 12:357–360. https://doi.org/10.1038/nmeth.3317

45. Liao Y, Smyth GK, Shi W (2014) Sequence analysis featureCounts : an efficient general purpose program for assigning sequence reads to genomic features. 30:923–930. https://doi.org/10.1093/bioinformatics/btt656

46. Love MI, Anders S, Huber W (2014) Differential analysis of count data – the DESeq2 package

47. Yang Y, Han L, Yuan Y, et al (2014) Gene co-expression network analysis reveals common system-level properties of prognostic genes across cancer types. Nat Commun 5:1–9. https://doi.org/10.1038/ncomms4231

48. Zhang B, Horvath S (2005) A general framework for weighted gene co-expression network analysis. Stat Appl Genet Mol Biol 4:. https://doi.org/10.2202/1544-6115.1128

49. Dennis G, Sherman BT, Hosack DA, et al (2003) DAVID: Database for Annotation, Visualization, and Integrated Discovery. Genome Biol 4:. https://doi.org/10.1186/gb-2003-4-9-r60

50. Szklarczyk D, Gable AL, Lyon D, et al (2019) STRING v11: Protein-protein association networks with increased coverage, supporting functional discovery in genome-wide experimental datasets. Nucleic Acids Res 47:D607–D613. https://doi.org/10.1093/nar/gky1131

51. Zhou G, Soufan O, Ewald J, et al (2019) NetworkAnalyst 3.0: A visual analytics platform for comprehensive gene expression profiling and meta-analysis. Nucleic Acids Res 47:W234–W241. https://doi.org/10.1093/nar/gkz240

52. Emmert-Streib F, Dehmer M, Haibe-Kains B (2014) Gene regulatory networks and their applications: Understanding biological and medical problems in terms of networks. Front Cell Dev Biol 2:1–7. https://doi.org/10.3389/fcell.2014.00038

53. Weirauch MT, Yang A, Albu M, et al (2014) Determination and inference of eukaryotic transcription factor sequence specificity. Cell 158:1431–1443. https://doi.org/10.1016/j.cell.2014.08.009

54. Paul Shannon 1, Andrew Markiel 1, Owen Ozier, 2 Nitin S. Baliga, 1 Jonathan T. Wang, 2 Daniel Ramage 2, et al (1971) Cytoscape: A Software Environment for Integrated Models. Genome Res 13:426. https://doi.org/10.1101/gr.1239303.metabolite

55. Byers LA, Diao L, Wang J, et al (2013) An epithelial-mesenchymal transition gene signature predicts resistance to EGFR and PI3K inhibitors and identifies Axl as a therapeutic target for overcoming EGFR inhibitor resistance. Clin Cancer Res 19:279–290. https://doi.org/10.1158/1078-0432.CCR-12-1558

56. Guo CC, Majewski T, Zhang L, et al (2019) HHS Public Access. 27:1781–1793. https://doi.org/10.1016/j.celrep.2019.04.048.Dysregulation

57. Tan TZ, Miow QH, Miki Y, et al (2014) Epithelial‐mesenchymal transition spectrum quantification and its efficacy in deciphering survival and drug responses of cancer patients. EMBO Mol Med 6:1279–1293. https://doi.org/10.15252/emmm.201404208

