## Supplementary materials for "Integrative systems biology approach identified crucial genes and transcription factors associated with gallbladder cancer pathogenesis"

### Supplementary Fig 1

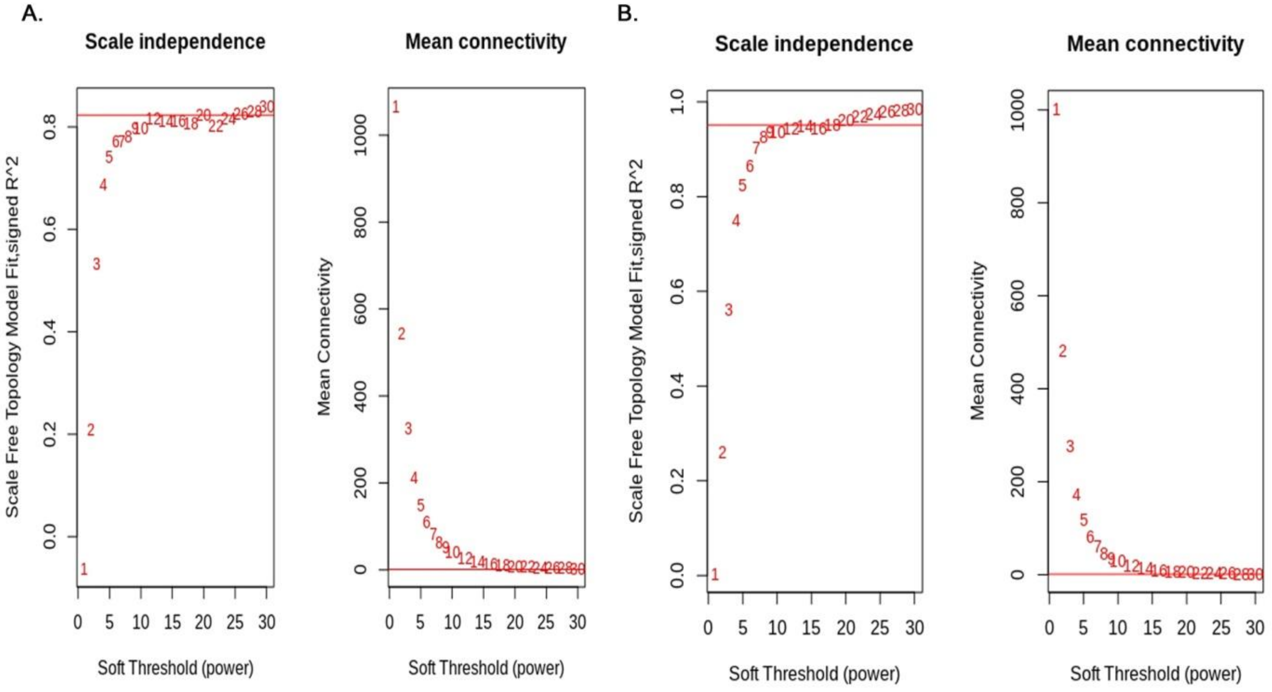


### Supplementary Table 1

| Module | medianRank.pres | Zsummary.pres |
|---|---|---|
| black | 16 | 2.2 |
| brown | 4 | 27 |
| cyan | 10 | 4.1 |
| darkgreen | 2 | 6.4 |
| darkgrey | 10 | 2.6 |
| darkorange | 7 | 2.6 |
| darkred | 5 | 5.3 |
| darkturquoise | 13 | 2.5 |
| gold | 12 | 24 |
| green | 17 | 2.1 |
| greenyellow | 8 | 4.9 |
| grey60 | 19 | 0.86 |
| lightgreen | 3 | 6.4 |
| lightyellow | 13 | 3.5 |
| magenta | 16 | 3 |
| midnightblue | 10 | 3.8 |
| orange | 19 | -0.12 |
| purple | 1 | 19 |
| royalblue | 8 | 3.7 |
| salmon | 20 | 1.4 |
| tan | 18 | 1.1 |
| white | 12 | 2.7 |

### Supplementary Table 2

| Modules | medianRank.pres | Zsummary.pres |
|---|---:|---:|
| blue | 3 | 29 |
| cyan | 11 | 4.2 |
| darkgreen | 9 | 2.8 |
| darkgrey | 4 | 3.6 |
| darkorange | 12 | 3.7 |
| darkred | 13 | 2.2 |
| darkturquoise | 6 | 4.1 |
| gold | 11 | 25 |
| green | 1 | 12 |
| greenyellow | 12 | 3.6 |
| grey | 20 | -0.49 |
| lightcyan | 16 | 6.3 |
| lightyellow | 13 | 2.3 |
| midnightblue | 16 | 0.91 |
| orange | 5 | 3.2 |
| pink | 13 | 3.5 |
| red | 14 | 5.1 |
| royalblue | 16 | 1.2 |
| tan | 16 | 2.3 |
| turquoise | 3 | 15 |

### Supplementary Table 3

| Sample name | GS76 | MLR | KS |
|---|---|---:|---:|
| GSM4146148 | 6.92582495614312 | 0.922485283 | -0.419032672 |
| GSM4146149 | -15.4277105379992 | 1.038697965 | 0.272364103 |
| GSM4146150 | -2.78710068072522 | 0.956055384 | -0.085746897 |
| GSM4146151 | 3.08850458758825 | 0.939545828 | -0.253673848 |
| GSM4146152 | 3.54856723716013 | 0.940623403 | -0.276644314 |
| GSM4146153 | -11.2641449810022 | 0.962427526 | 0.121129976 |
| GSM4146154 | 7.92566592889314 | 0.931538294 | -0.382222856 |
| GSM4146155 | 4.85039097543316 | 0.924028816 | -0.223997717 |
| GSM4146156 | 5.34065727847344 | 0.920960796 | -0.30960194 |
| GSM4146157 | 5.83634613093576 | 0.935251049 | -0.229847339 |
| GSM4146158 | 11.6216471377933 | 0.887363848 | -0.330860322 |
| GSM4146159 | -19.0938301382121 | 1.06273886 | 0.357968326 |
| GSM4146160 | 12.4611940094735 | 0.932319314 | -0.486517335 |
| GSM4146161 | 6.64607743670175 | 0.942107475 | -0.256955343 |
| GSM4146162 | 11.4271108425853 | 0.903626614 | -0.482950492 |
| GSM4146163 | -18.4380535030102 | 1.046387984 | 0.277928378 |
| GSM4146164 | 11.1470869283106 | 0.903418062 | -0.58952775 |
| GSM4146165 | 3.95593866468601 | 0.925905771 | -0.212726495 |
| GSM4146166 | -9.47737699957706 | 1.038349726 | 0.130831788 |
| GSM4146167 | -18.2867952736515 | 1.038623957 | 0.247253531 |
